# Social hierarchy position in female mice is associated with plasma corticosterone levels and hypothalamic gene expression

**DOI:** 10.1101/529131

**Authors:** Cait M. Williamson, Won Lee, Alexandra R. Decasien, Alesi Lanham, Russell D. Romeo, James P. Curley

**Affiliations:** Department of Psychology, Columbia University, New York, NY, 10027, USA; Department of Anthropology, New York University, New York, NY 10003, USA; New York Consortium in Evolutionary Primatology, New York, NY, 10024, USA; Department of Psychology, Barnard College, New York, NY, 10027, USA; Department of Psychology, University of Texas at Austin, Austin, Texas, 78712, USA

## Abstract

Social hierarchies emerge when animals compete for access to resources such as food, mates or physical space. Wild and laboratory male mice have been shown to develop linear hierarchies, however, less is known regarding whether female mice have sufficient intrasexual competition to establish significant social dominance relationships. In this study, we examined whether groups of outbred CD-1 virgin female mice housed in a large vivaria formed social hierarchies. We show that females use fighting, chasing and mounting behaviors to rapidly establish highly directionally consistent social relationships. Notably, these female hierarchies are less linear, steep and despotic compared to male hierarchies. Female estrus state was not found to have a significant effect on aggressive behavior, though dominant females had elongated estrus cycles (due to increased time in estrus) compared to subordinate females. Plasma estradiol levels were equivalent between dominant and subordinate females. Subordinate females had significantly higher levels of basal corticosterone compared to dominant females. Analyses of gene expression in the ventromedial hypothalamus indicated that subordinate females have elevated ERα, ERβ and OTR mRNA compared to dominant females. This study provides a methodological framework for the study of the neuroendocrine basis of female social aggression and dominance in laboratory mice.

## Introduction

The contextual and neurobiological factors that influence male intrasexual aggression and social dominance have been well-studied across species^1–6^. Conversely, female aggression and social dominance have been relatively understudied, with most work focused on maternal aggression expressed by females when they are pregnant or during the early postpartum period where the behavioral focus is on maternal defense of offspring^7,8^. Few studies have investigated the contextual and neurobiological factors that influence female-female aggression outside of reproduction in rodents^9,10^.

Social hierarchies are likely to emerge whenever there is competition between individuals for resources such as food, water, territory or access to mates^11^. The more intense this competition is, the more likely it is that a highly linear social hierarchy will develop. In mammals, male social hierarchies are common as inter-sexual competition is typically dramatically higher in males compared to females, though there are notable exceptions such as hyenas where females have high levels of intra-sexual conflict and form strong female hierarchies^11^. Female hierarchies have also been observed in other species that have female intrasexual competition for access to resources including degus^12^, bison^13^, caribou^14^, red deer^15^, vervet monkeys^16^, and chimpanzees^17^. Less is known about the formation of social hierarchies in female wild mice, though some population studies suggest that females do generate some form of social hierarchy with dominant aggressive females establishing territories and subordinate females being unable to do so when population sizes increase^18,19^. Conversely, when population density is very low it appears that female wild mice have relatively little intra-sexual competition and do not form hierarchies^20^. Female-female aggression also appears to be low if females have social experience with each other prior to the intra-sexual competition^21,22^. Conversely, small groups of female laboratory mice can establish social ranks based on home cage social interactions^23^ or their performance in the tube-test^24,25^.

Previously, we have explored the complex group dynamics and neurobiology of male social hierarchies, demonstrating that male outbred CD-1 mice living in groups of up to 30 individuals will form highly linear social hierarchies when living in a large laboratory-based vivarium^5,26,27^. As relatively little is known about whether large groups of non-reproductively active female mice will form social hierarchies in the laboratory, we aimed to explore this question by housing groups of 12 virgin outbred CD-1 female mice in large vivaria. One historical reason why female behavior is so vastly understudied in comparison to male behavior in laboratory rodents is due to concerns that female behavior is more variable than males due to fluctuations in steroid hormone levels across the female estrus cycle. Indeed, estrus state has been shown to influence many behavioral states including anxiety-like behavior and exploration^28^, motivation, addiction^29^ and fear^30^. In rodents, some species also show variation in aggressive behavior across the estrous cycle^31^. Female California deer mice^32^, rats^33,34^ and hamsters^35–38^ are less likely to show aggressive behavior during estrus than diestrus, although other studies have found no effect of estrus state on aggressive behavior^39–42^. There is also mixed evidence for estrous effects on aggression in female house mice.^43^ ^44^. Given the potential significance of estrous state on female dominance and subordinate behaviors, we examined whether the estrous state of females is associated with the frequency of aggressive behavior within social hierarchies.

The neurobiological basis of female intrasexual aggression among non-reproductive females is receiving increased attention though much less is still known compared to male intrasexual aggression^9,10^. As in males, brain regions in the social behavior network (medial amygdala (meA), bed nucleus of the stria terminalis (BNST), lateral septum (LS), medial preoptic area (mPOA), anterior hypothalamus (AH), ventromedial hypothalamus (VMH) and periaqueductal grey (PAG)) as well as the mesocorticolimbic dopamine pathway have been found to form the basis of the neural circuit regulating aggression, though there are some important sex differences^10^. In particular, it is well-established that the VMH is a key modulator of aggression in non-reproductive female rodents^9^. Further, estradiol, the major estrogen steroid hormone, has been primarily associated with promoting aggressive behaviors in females^45–47^. Estradiol acts to alter the expression of gene products in the hypothalamus, including progesterone receptors (PR), oxytocin receptors (OTR), opioid receptors, and gonadotropin-releasing hormone (GnRH), all of which are known to regulate female social behaviors including social recognition, memory and aggression^48^.

The current study used an established paradigm developed in our lab applied to the study of male social hierarchy dynamics to study female social hierarchy behavior and begin to disentangle underlying neurobiological and neuroendocrine mechanisms. We investigated the hierarchical structure of eight groups of twelve females as well as plasma corticosterone and plasma estradiol concentrations for dominant and subordinate mice within these hierarchies. In male mice, social stressors such as social defeat and social stability are known to lead to increases in basal levels of corticosterone but findings from female mice are more variable^49,50^. We have previously found that subordinate male mice have elevated corticosterone levels compared to dominant male mice but only if the dominant males are highly despotic. We predicted that female subordinate mice may also show higher levels of basal corticosterone compared to dominant females, although previous studies have not found a consistent relationship between social dominance and plasma corticosterone in female mice^23,51^. We further examined gene expression differences between dominant and subordinate individuals in the VMH and mPOA of the hypothalamus across six genes known to modulate various aspects of social behavior and moderated by the action of estrogen: ERα, ERβ, PR, OTR, OPRM1, and GnRH. The aim of this work was to establish a feasible methodology for the study of female aggression and dominance, as well as their neurobiological and neuroendocrine mechanisms, outside of the reproductive period in laboratory mice.

## Results

### Hierarchy measures and organization

Sociomatrices of win-loss data for all female groups as well as the emergence of individual dominance ranks over time are presented in Figure 1 **and** supplemental **Figure S1**. Summary statistics of several aspects of the hierarchical structure of each group are provided in Table 1. We found that seven out of the eight social groups formed a significantly linear and steep social hierarchy as measured by modified Landau’s *h’*, triangle transitivity and steepness. All eight social groups had significantly high directional consistency of agonistic behavior, indicating that the majority of wins were directed from more dominant to more subordinate individuals. Female groups had relatively low despotism values, indicating that alpha females were not exerting complete dominance over all other females. This interpretation was confirmed by moderate Gini Coefficient values for wins, demonstrating that the number of wins made by dominant females was fairly evenly distributed between the top ranked females. Initial body weight measured at the beginning of group housing was not related to final dominance rank in any group (Spearman Rank correlation tests: all p>0.12).

**Table 1.**
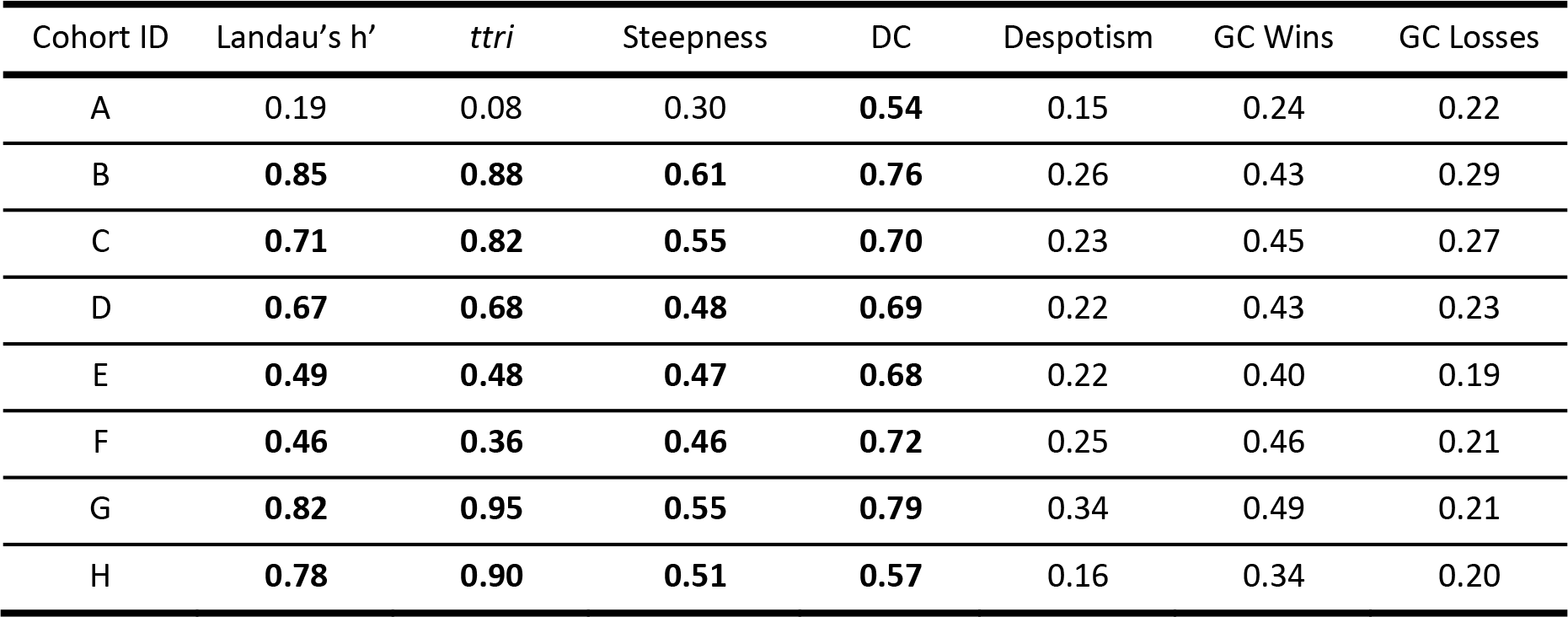
Social hierarchy measures for each cohort (A-H). Females have significantly linear hierarchies (as measured by Landau’s Modified h’ value and triangle transitivity (ttri)) as well as significantly steep hierarchies (steepness). Aggressive behavior is also significantly directionally consistent (DC). Significant values are bolded. Despotism and the Gini-Coefficient of Wins and Losses measure how evenly distributed aggressive behavior is across ranks. Female hierarchies are not highly despotic.

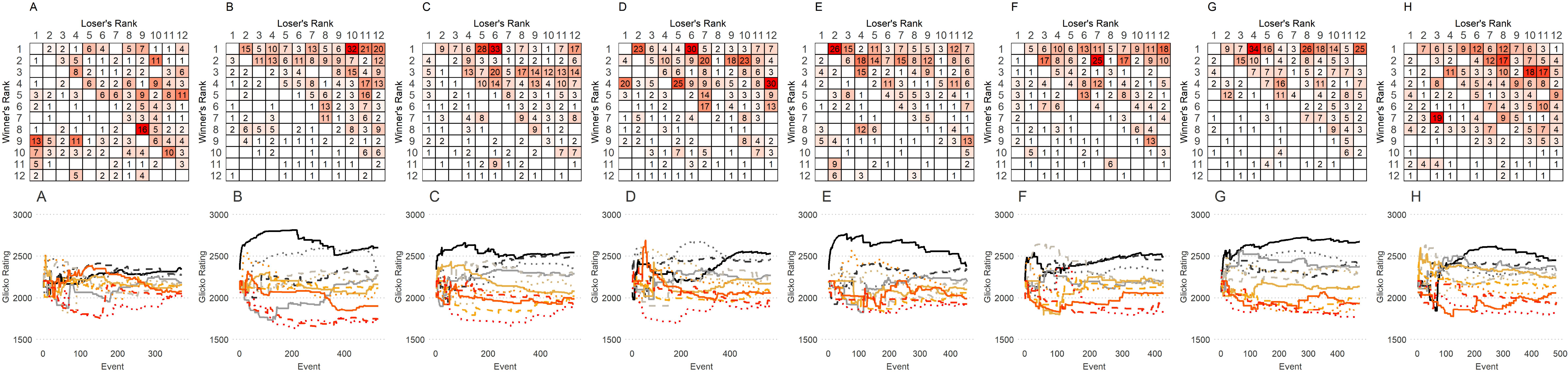

### Emergence of hierarchies over time

Despite having no previous social experience with each other, 4 of 8 social groups had a significantly linear hierarchy by the end of Day 1 that continued throughout the 14-day observation period (Figure 2). Two further social groups were significantly linear by the end of Day 2 and thereafter. One cohort did not have a stable linear hierarchy until Day 9, although this hierarchy did show significant triangle transitivity on Days 1 and 4 suggesting some linearity in the first week of co-housing. The social group that did not have a significant hierarchy by the end of the observation period did have some linear organization, having a significantly linear hierarchy on Day 3.

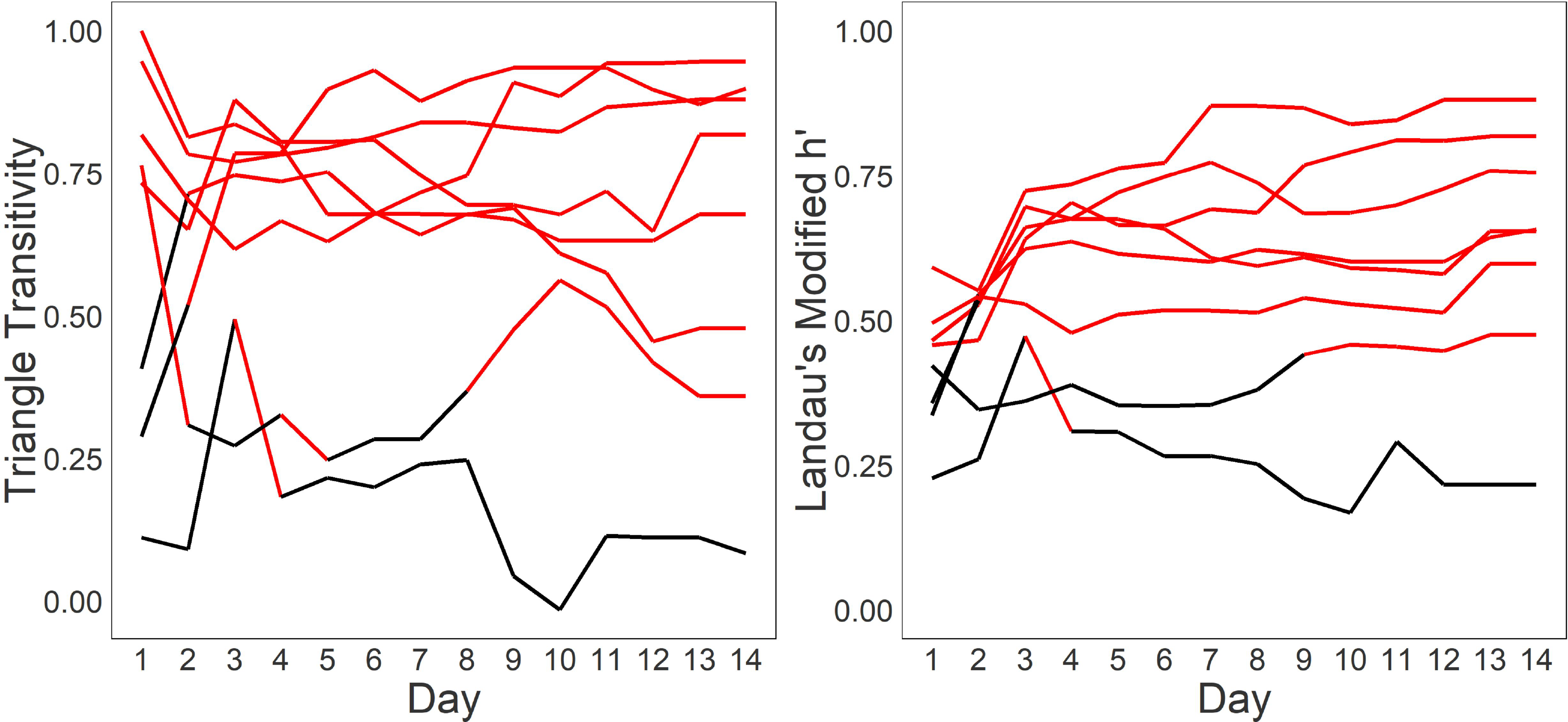

### Sex differences in mouse social hierarchies

Female social hierarchies were significantly different from male social hierarchies in several aspects of their dominance organization (see Methods about male comparison group). Female social hierarchies were significantly less linear by triangle transitivity (W=14, p<0.05) and had lower directional consistency (W=0, p<0.001) than male social hierarchies (Figure 3). The distribution of power was more even in females than in males. Alpha females were significantly less despotic than alpha males (W = 1, p<0.001), and the Gini coefficient of wins (W=0, p<0.001) was significantly lower in female hierarchies than male hierarchies. There was no difference in the steepness of hierarchies (W=31, p=0.460), the Gini coefficient of losses (W=32, p=0.515.) or Landau’s modified h’ value (W=23, p=0.146).

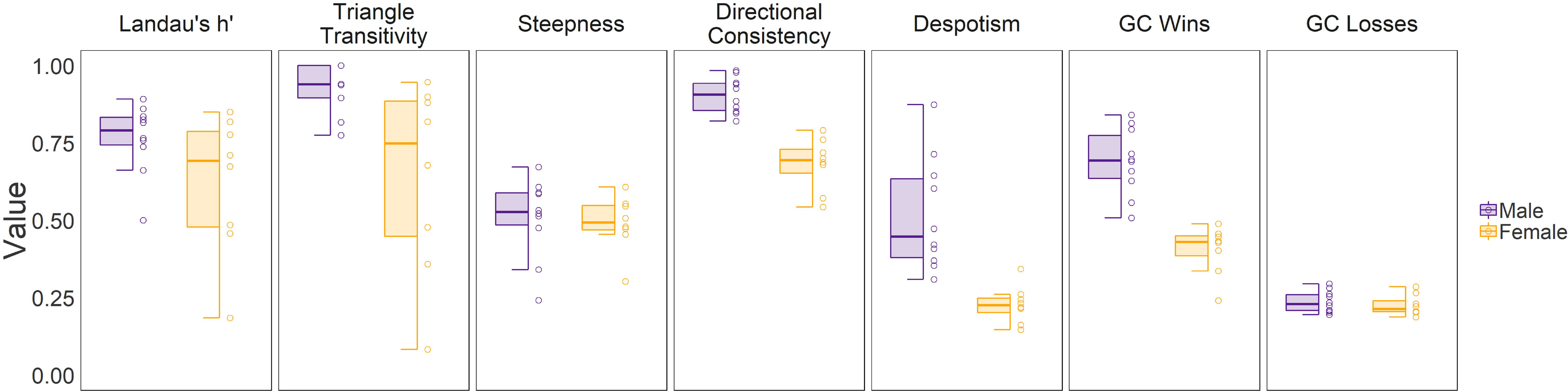

### Frequency of each agonistic behavior over the group housing period

The hourly rate of each agonistic behavior is shown in **Figure S2**. Fighting and chasing were observed at significantly higher rates than mounting. The rate of fighting behavior showed a significant decrease by day over the group housing period (b_day_= −0.62 [−0.89, −0.36]) while those of chasing and mounting behaviors did not show significant changes by day (chasing: b_day_= 0.08 [−0.29, 0.44]; mounting: b_day_= 0.10 [−0.03, 0.23]).

### Relationships between mounting, chasing and fighting

To determine if the directionality of fighting, chasing and mounting was consistent between individuals we performed QAP correlation tests on the fighting, chasing and mounting sociomatrices (**Figure S3)**. Fighting and chasing sociomatrices were highly correlated across all cohorts (data presented as median [IQR] across all eight cohorts: r=0.79 [0.75, 0.80], all p<0.001). Chasing and mounting sociomatrices were also significantly correlated across all cohorts (r=0.44 [0.38, 0.54], all p<0.025). Fighting and mounting sociomatrices were correlated for 7/8 cohorts (r=0.37[. 35,.39], all p<.025 for significant correlations). See supplemental **Figure S3** for individual sociomatrices.

### Directional consistency of each agonistic behavior

All groups showed highly significant directionally consistent behavior for all behaviors (all p<0.001). Notably, the most directionally consistent behavior was mounting behavior (median [IQR] = 0.87 [0.83, 0.93]), which was more consistent than chasing (0.75 [0.70, 0.77]) and significantly more consistent than fighting (0.72 [0.69, 0.74]) (**Figure S4A**; Friedman’s test X^2^=7,df=2, p=0.03; post-hoc test mounting vs fighting p= 0.033, mounting vs chasing p= 0.112).

### Frequency of each agonistic behavior across individuals and ranks

As expected, the Gini coefficient for the total number of attacks and chases made by the females in each group was moderately high, indicating that a few females are responsible for a disproportionate number of these aggressive acts (**Figure S4B and S4C**). Unexpectedly, the Gini coefficient for mounting other females was even higher than for fighting or chasing across cohorts (Friedman Test: X^2^ = 12, df=2, p=0.002; post hoc tests p<0.01). This analysis demonstrates that in each social group a small number of females are responsible for a very large proportion of all mounting acts. The Gini coefficient for the total number of attacks and chases received by each female in each social group was moderately low. This finding indicates that most females are the recipients of fights and chases and these events are relatively evenly distributed across females in each social group. Again, it was unexpected that the Gini coefficient of mounts received was significantly higher than that for receiving the other two behaviors (Friedman Test: X^2^ = 9.2, df=2, p=0.01; post hoc tests p<0.05). This finding indicates that being mounted is far more unequally distributed across females in each group than being chased or being attacked: certain females in each social group are the targets for a disproportionately high number of mounts from other females.

We further examined if there is an effect of social rank on the hourly occurrence rate of each agonistic behavior either given or received. Fighting and chasing, but not mounting, were exhibited at significantly higher hourly rates by more dominant females (Figure 4, fighting given: b_rank_= −1.11 [−1.44, −0.81]; chasing given: b_rank_= −1.13 [−1.46, −0.85]; mounting given: b_rank_= −0.25 [−0.61, 0.12]). Similarly, there were significant effects of social rank on the hourly rate of receiving fighting and chasing but not mounting, showing more subordinate females received more fighting and chasing behaviors (fighting given: b_rank_= - 0.38 [0.19, 0.58]; chasing given: b_rank_= 0.47 [0.25, 0.71]; mounting given: b_rank_= 0.10 [−0.20, 0.42]). Although animals of each rank differed in the absolute frequencies of each agonistic behavior used, there was no effect of rank on the proportion of each behavior used (**Figure S5**). That is, all animals use each behavior proportionally equivalently.

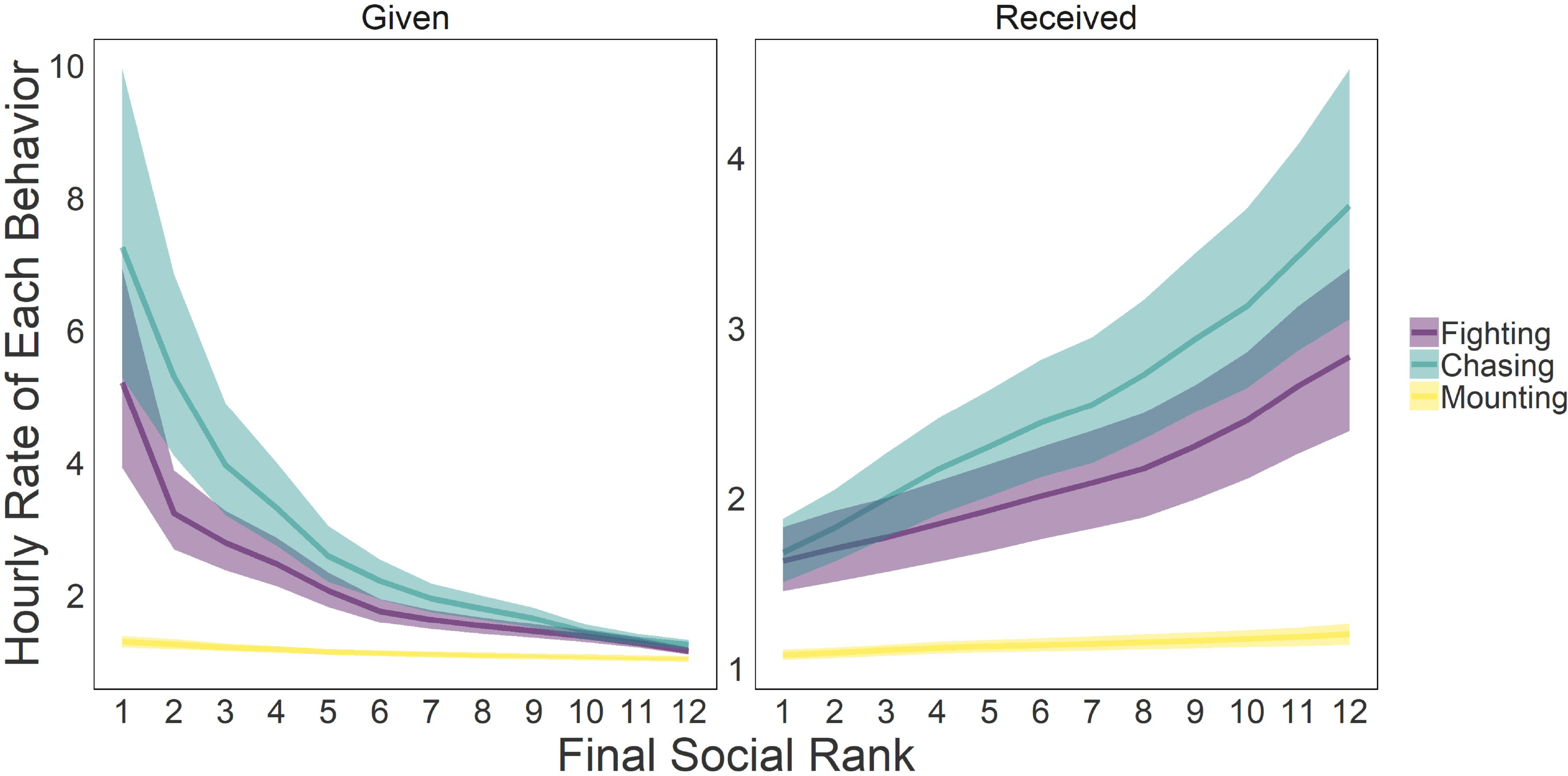

### Estrus cycle and female behavior

The median (+/− IQR) proportion of time that individuals were in each estrous state were - estrus: 33.3% [25.8%, 46.7%]; proestrus: 21.4% [13.3%, 36.4%], diestrus: 14.3% [0.08%, 23.1%], metestrus: 21.4% [14.3%, 28.6%] (Figures. 5A and 5B). Female mice with higher social rank were more likely to be in estrus compared to metestrus or proestrus (log-odds of being in estrus vs. metestrus: 0.48 [0.02, 0.95]; estrus vs. proestrus: 0.36 [018, 1.57]). Mice did not significantly differ by social rank in the likelihood to be in estrus compared to diestrus (estrus vs. diestrus: 0.50 [−0.03, 1.06]), or between other states (metestrus vs. proestrus: 0.32 [−0.36, 1.05]; metestrus vs. diestrus: 0.02 [−0.57, 0.58]; proestrus vs. diestrus: −0.22 [−0.78, 0.35]).

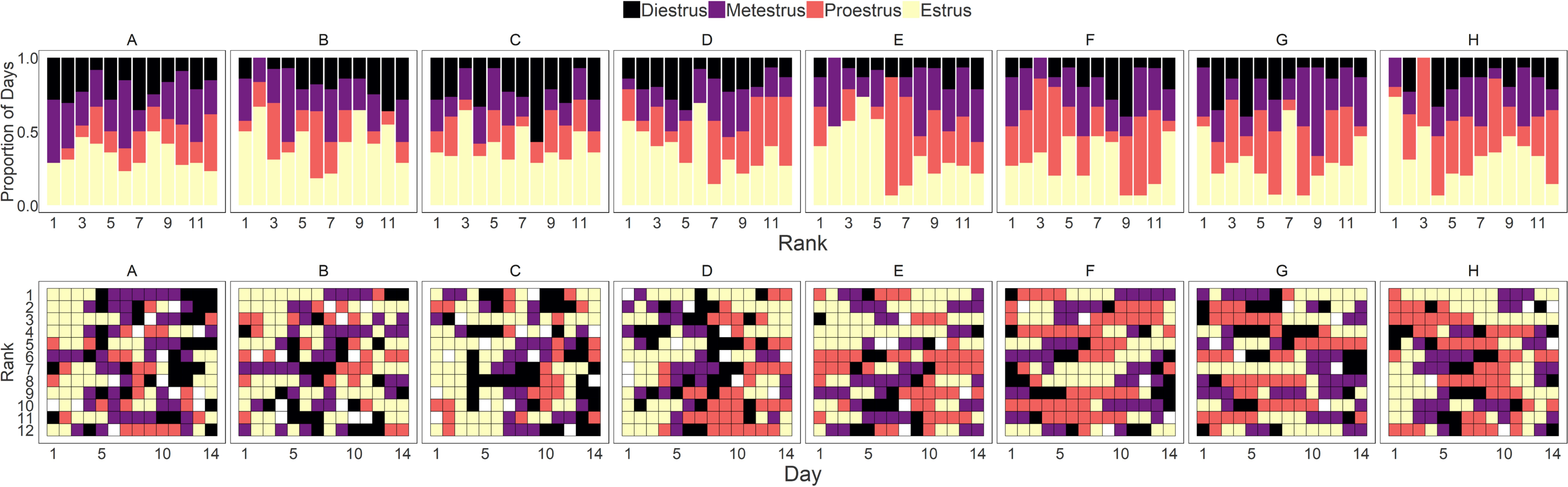

Overall there was no large effect of daily estrus state on the hourly rate of giving or receiving aggression. There was small effect that mice showed a higher hourly rate of giving aggression when in metestrus compared to proestrus (b_metestrus-proestrus_: 0.19 [0.005, 0.35]). We further examined the effect of daily estrus state on the hourly rate of giving or receiving each agonistic behavior individually (fighting, chasing, mounting). For the rate of giving chases, mice had a higher rate when they are in metestrus compared to diestrus and proestrus (b_metestrus-diestrus_: 0.22 [0.03, 0.40]; b_metestrus-proestrus_: 0.19 [0.01, 0.36]). Compared to during diestrus, mice received higher rates of mounting when they were in estrus or metestrus (b_estrus-diestrus_: 0.31 [0.09, 0.53]; b_metestrus-diestrus_: 0.33 [0.08, 0.58]). The hourly rates of other behaviors given or received did not differ across different estrus states.

### Plasma corticosterone and estradiol levels

Plasma corticosterone levels were found to be significantly higher for subordinate individuals as compared to dominant individuals (Figure 6A, b_subordinate-dominant_: 153.9 ng/ul [107.6, 200.0]). We did not find an effect of estrus cycle state on corticosterone level. There was no effect of dominant-subordinate status on plasma estradiol levels (Figure 6B, b_subordinate-dominant_: −2.97 pg/ul [−8.03, 1.99]). Mice that were in diestrus state on the day of blood collection had significantly higher estradiol levels compared to those that were in metestrus or proestrus (b_diestrus-metestrus_: 6.79 pg/ul [0.29, 13.2]; b_diestrus-proestrus_: 10.1 pg/ul [3.07, 17.5]). There was no effect of estrus state on the estradiol levels among other states.

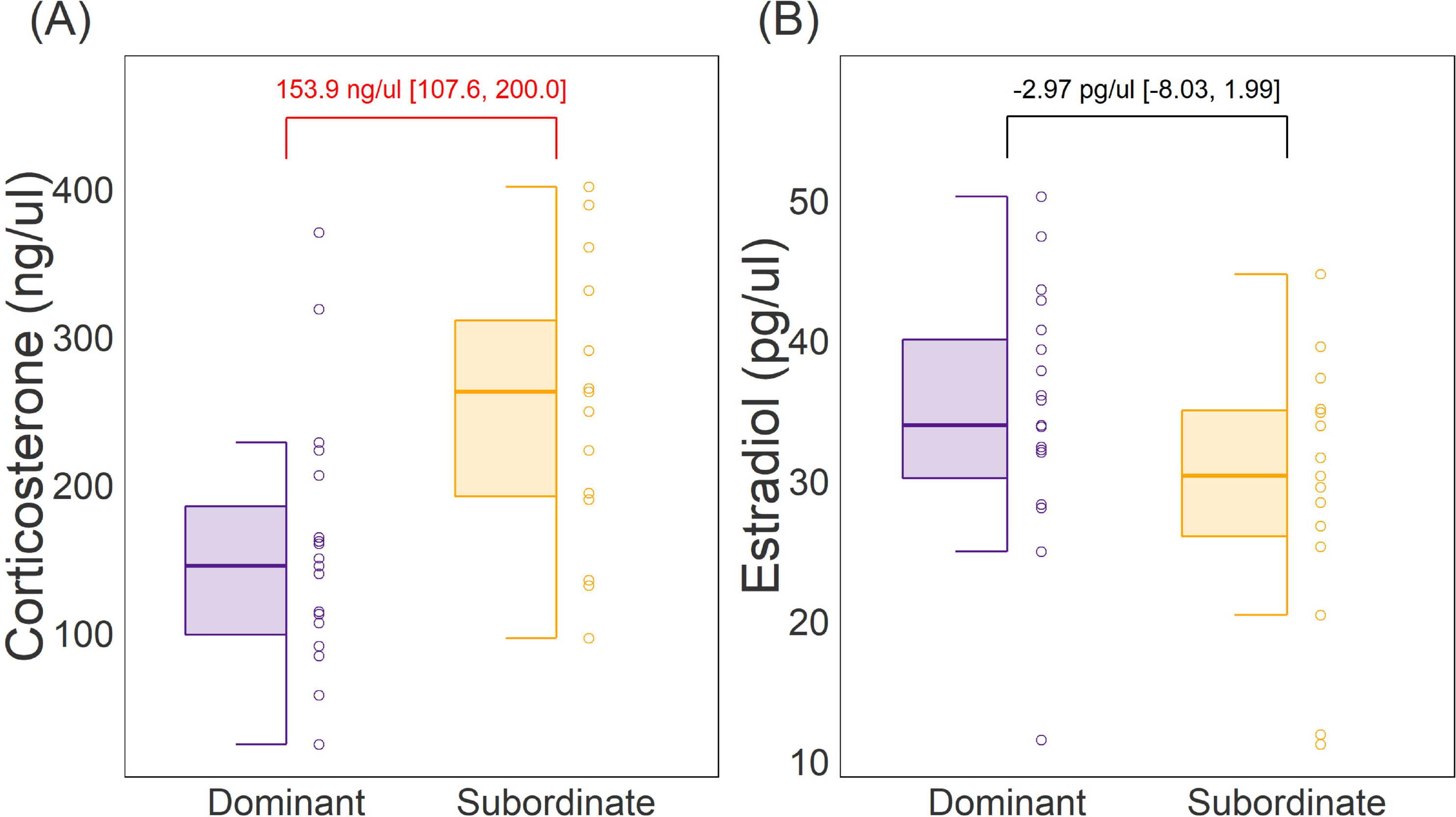

### Gene expression in the VMH and the mPOA

In the VMH, there was small but significant effect of dominant-subordinate status on the levels of expression in the VMH of ERα, ERβ, and OTR genes (Figure 7), with subordinate mice showing higher expression than dominants. There were no significant differences in expression levels of OPRM1 and PR genes in the VMH. There were no significant differences between dominant and subordinate individuals in the expression any of the genes examined in the mPOA.

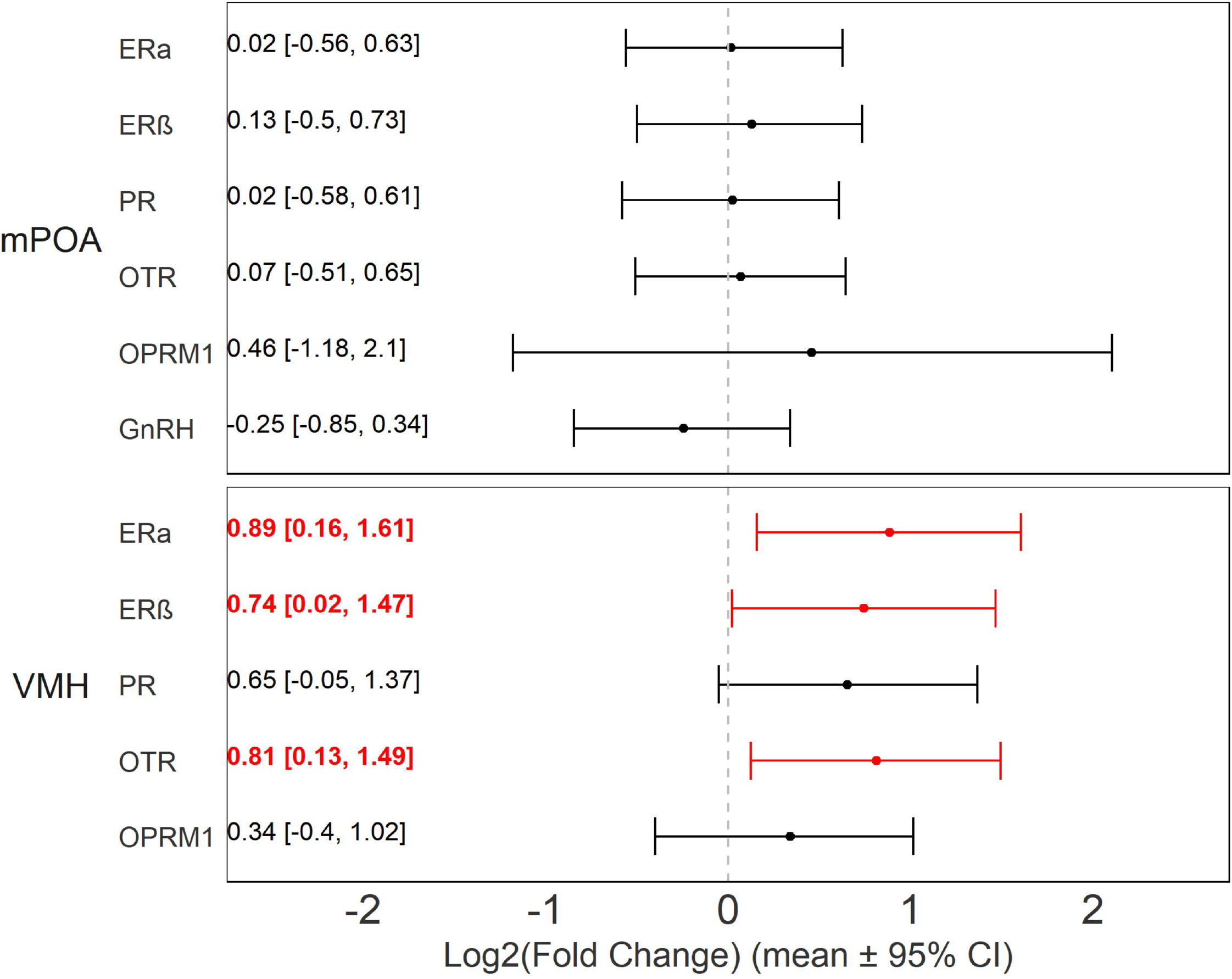

## Discussion

Here we show that female mice living in social groups of up to 12 females are capable of forming significantly linear dominance hierarchies that are stable for up to 14 days. Seven out of eight female social groups had significantly linear hierarchies, as measured by Landau’s h-value and triangle transitivity, and significantly steep hierarchies as measured by the relative differences in David’s scores. All eight female social groups had significantly high directional consistency of agonistic behavior demonstrating that dominant individuals more often won competitive interactions against subordinate individuals. Similar to males, female mammals can form stable dominant-subordinate relationships whenever there is intrasexual competition for resources^23,24,52–54^. In our study of virgin female mice, there is no intrasexual competition for access to mates or food and water, however, female mice may compete for access to preferred areas of the vivaria (such as nestboxes) or bedding. Our results also extend the findings from studies of round-robin tube-tests conducted in groups of 5 or 8 female mice that report that females were capable of forming linear social hierarchies based on wins and losses in that competitive exclusion test^24,25^. These findings are also consistent with studies of wild mice that have shown that females will form territorially based social hierarchies when living in large environments with relatively high population densities^18,19^. It appears that there may be a critical population density limit required for there to be sufficient intrasexual competition for females to engage in sufficient aggression to establish hierarchies, as one recent study found that groups of six females living in enclosures of 7m^2^ did not form social hierarchies^20^.

Our results demonstrate that females use fighting, chasing and mounting behaviors to establish and maintain dominance relationships. The highest rates of fighting were observed during the establishment of hierarchies (which typically occurred rapidly within the first two days), and the rate of fighting significantly decreased thereafter. The rates of chasing and mounting did not decrease across the housing period. Unsurprisingly, given the existence of a linear hierarchy, higher ranked individuals exhibited higher rates of fighting and chasing than lower ranks. Similarly, lower-ranked individuals received higher amounts of each of these behaviors. With respect to mounting behavior, we did not observe a significant overall effect of rank on the rate of giving or receiving mounting. Indeed, there has been some controversy as to the proximal function of female-female mounting in rodents and whether it should be considered to be a dominance behavior, a sexual behavior, a masculinized behavior or something else^55–57^. Our data suggest that mounting is being used by some females as a dominance behavior but not consistently by all individuals. That is, females are more likely to mount females subordinate to themselves but not all females engage in this behavior. We observed that the group sociomatrices for fighting, chasing and mounting behaviors given and received all highly correlated with each other, indicating that the direction and magnitude of these behaviors within each social relationship was consistent. Secondly, we observed that the directional consistency of mounting behavior (0.87) was even stronger than the directional consistency for fighting (0.72) and chasing (0.75). This finding indicates that mounting behavior consistently occurred 87% of the time in the direction of more dominant to more subordinate females. We also noted that females of all ranks, despite having different absolute rates of each behavior, used each of the three behaviors in roughly equal proportions-suggesting that higher- or lower-ranked females do not preferentially utilize mounting behavior over the other two as a dominance behavior. Additionally, the Gini coefficient of behavior given and received was significantly higher for mounting as compared to the other two agonistic behaviors. Thus, in each social group the majority of mounts given and received were by specific individuals, and these interactions occurred in a directionally consistent manner.

Female mouse hierarchies exhibit several differences when compared to male social hierarchies. Although both sexes produce linear hierarchies, those of females are less linear and have lower directional consistency than male hierarchies. Male hierarchies tend to be despotic, with one alpha male exhibiting the vast majority of all aggressive acts^5,26,27^. We did not observe such despotism in female hierarchies as evidenced by the lower despotism scores and lower Gini coefficient of wins. More dominant females in social groups tend to more equally distribute aggression towards other lower-ranked females. These sex differences may be rooted in the ancestral biology of the house mice. In the wild, there are fundamental differences in the reproductive strategies of males and females. Males have high levels of intersexual competition and have high reproductive skew^58^. Females generally exhibit higher parental investment in offspring and have reduced conflict over reproductive opportunities^54,59,60^. Indeed, even when given the opportunity to compete for males, female mice do not necessarily increase their rates of aggression^20^. Across mammals, female adaptations for intrasexual competition can involve more subtle behaviors such as low-level persistent aggression instead of overt displays of physical aggression^54^. Our data suggest that in laboratory mice there persists low-level aggression that results in females forming stable social hierarchies even with unlimited access to food, water, space and nesting material and no access to males.

We found that the estrous state of each female did not have a large effect on the likelihood of females giving or receiving aggression. Females in all states were equally likely to bite or be bitten. However, we did observe a very small effect for females in metestrus to have a higher rate of chasing than those females in proestrus or diestrus. Previous work in rodents has suggested that estrus state can influence the propensity of females to engage in intrasexual aggression although these effects are inconsistent and appear to be highly influenced by many other contextual factors. For instance, female mice have been reported to show higher aggression in the resident-intruder test during metestrus and proestrus compared to estrus and distrus^43^. Similarly, female California mice show their highest aggression during diestrus compared to proestrus and estrus^32^. Such a decline in aggression during estrus has been suggested to facilitate mating, yet female mice in estrus have been shown to be capable of exhibiting aggression and forming dominant-subordinate relationships^61^. Conversely, no changes in female-female aggression across the estrous cycle have also been observed in rats and hamsters^41,42^. Although there was no effect of estrous state on the likelihood to mount other females, we did find that female mice in estrus or metestrus received significantly more mounts than those females in diestrus. This finding is somewhat consistent with evidence from several species including baboons^62^, squirrel monkeys^63^, hanuman langurs^64^, rabbits^65^ and rats^56^ that mounted females tend to be subordinates in estrus.

We observed that group-housed females had extended estrous cycles with prolonged periods of diestrus and estrus. The mouse estrus cycle is regulated by luteinizing hormone and follicle-stimulating hormone released from the pituitary in response to gonadotrophin releasing hormone released from the hypothalamus, which is in turn under control from estrogen and progesterone. The duration of this cycle is usually 4-6 days for females housed in isolation or in pairs, but can be much longer for group-housed females – an effect known as the Lee-Boot effect^66–69^. For instance, female mice living in groups of 8 have been shown to have estrous cycles lasting up to 14 days^66^ and females in groups of 30 have estrus cycles up to 40 days in duration^69^. These cycles are typically extended due to longer periods of time spent in diestrus, but they may also become disrupted. Following exposure to males, females quickly enter estrus and become sexually receptive^69^. Our results are largely congruent with these earlier studies in that we do observe extended estrous cycles in group-housed animals. Notably, many females in our study were in estrous for prolonged periods of time, and this effect was larger in dominant females compared to subordinate females. It is possible that this is an adaptive mechanism by which dominant females ensure more or earlier mating opportunities than subordinate females, however it is unclear as to what physiological processes underpin the extended length of estrous cycles we observed.

We found no significant differences between dominant and subordinate mice in plasma estradiol. Peripheral estrogens are known to promote aggression in both males and females^10,70,71^, and therefore we predicted there would be higher estradiol levels in dominant females. However, our hormone samples were taken at the end of group housing on Day 14 at a time when the social hierarchies had stabilized and aggression was at a low level. Further, estrogens can have many, sometimes opposite, effects, depending on where in the brain and on what receptors they are acting^72,73^. It is therefore perhaps not surprising that estradiol levels found in plasma at one time point do not differ between dominant and subordinate individuals.

Conversely, we did find that subordinate females had significantly higher levels of plasma corticosterone than dominant females. This effect was moderately large and much larger than the effect that we had previously observed in male hierarchies, where significant differences in plasma corticosterone levels between dominants and subordinates only occur in highly despotic hierarchies^27^. In females, all hierarchies were considered to be very low on the despotism scale. These results are interesting in the context of the established literature on sex differences in stress responses. Several studies have shown that male mice show a much more robust physiological response to stressors such as social defeat, social instability and chronic stress compared to female mice^49,50^. However, these results are not always consistent, and sometimes females do show increases in corticosterone depending on the social context of the stressor^74–76^. It has also been reported that highly aggressive territorial females living in large groups have elevated corticosterone compared to non-aggressive females^51^, though one other study found results consistent to ours that subordinate female mice had higher corticosterone than dominant females in small groups of up to five albino mice^23^. Elevated levels of corticosterone in subordinates post-hierarchy formation have been shown to facilitate social memories for being socially subordinate in rats ^77,78^. The functional significance of the elevated corticosterone in subordinate females living in our relatively stable social housing remains to be addressed, but these data suggest the potential for studying female social hierarchies as a model of social stress.

The neurobiology of female intrasexual aggression has been relatively under-studied compared to that of male intrasexual aggression, although this area has started to receive increased attention^9,10^. Here we found that the expression of ERα, ERβ and OTR was moderately higher in subordinate female mice compared to dominant female mice in the VMH. The VMH is known to be a critical regulator of female aggression. Early studies in rats and hamsters demonstrated that lesions of the VMH lead to increased aggression by females^45,79^, whereas implants into the VMH with either estradiol or progesterone reduce aggression^42,80,81^. More recently in mice, estrogen-receptor expressing neuronal populations in the ventrolateral VMH have been found to become active when females bite versus mount other females^82^. This work builds on previous studies that demonstrated an important role for central estrogen receptors in regulating male and female aggression. ERα knockout male mice show reduced aggression towards other males^83^, whereas females lacking ERα expression are more aggressive to other females^84,85^. Loss of ERβ has also been associated with increased aggression in males^86,87^. Central administration of selective ERα agonists to female mice increases aggressive attacks in a resident-intruder paradigm^88^, while selective ERβ agonists treatment leads to an increase in non-aggressive social behaviors^89^. The role of each of these receptors in coordinating female aggressive behavior is clearly complex and context-dependent, but it is possible that our observed increased VMH expression in both ERα and ERβ in subordinate mice may underlie their inhibition of aggression. However, given the multitude of functions of these receptors in this region it is possible that these differences may be unrelated to aggression and associated with other social behaviors such as social recognition, learning and memory ^90^.

Likewise, oxytocin acting on oxytocin receptors has a range of effects on social behavior in females, and its roles in promoting or inhibiting aggression are highly contextually dependent. In non-lactating females oxytocin generally appears to inhibit aggression^10^. OT knockout mice show reduced aggression to each other^91^ and central or injections of OT into the MPOA or AH can reduce aggression^92,93^. These findings may be congruent with our finding that subordinate females have higher VMH OTR expression than dominant females. Notably, we did not observe any association between social status and mRNA levels of OTR or any other gene in the MPOA. Though some lesion studies in other rodents have reported a role for the MPOA in female aggressive behavior^94^, our results would suggest that plasticity in gene expression in response to an individual’s social status occurs primarily in the VMH.

## Conclusion

In the present study, we establish that outbred CD-1 female mice living in groups of 12 individuals are capable of forming significant linear hierarchies. These hierarchies are linear and directionally consistent, but less despotic than male social hierarchies. These hierarchies emerge quickly and are stable over 14 days and are relatively unaffected by the estrous cycle. All group-housed females also show an extended estrous cycle, and dominants spend longer in estrus than subordinate females. Subordinate females have significantly higher levels of plasma corticosterone than dominant females, suggesting that subordinate females are more susceptible to the social stress of group living than male mice. We also find that subordinate females have higher levels of ERα, ERβ, and OTR mRNA than dominant females in the VMH, suggesting that these genes in this region may facilitate in part the reduced aggression displayed by these females. This work furthers our understanding of group female social behavior, begins to explore sex differences between male and female social hierarchy formation and maintenance and provides evidence that the actions of estrogen may play a role in modulating female social hierarchy behavior.

## Methods

### Subjects and housing

A total of 96 female outbred CD1 mice were obtained from Charles River Laboratories at 7 weeks of age. Mice were housed in the animal facility in the Department of Psychology at Columbia University, with constant temperature (21-24°C) and humidity (30-50%). The room was kept on a 12/12 light/dark cycle, with white light (light cycle) on at 2400 hours and red lights (dark cycle) on at 1200 hours. All mice were uniquely marked by dying their fur with a blue, nontoxic animal marker (Stoelting Co.), enabling individuals to be identified throughout the study. These marks remain for up to 12 weeks and only require one application. All procedures were conducted with approval from the Columbia University Institutional Animal Care and Use Committee (IACUC Protocol No. AC-AAAP5405).

### Social behavior observations

Following arrival at the animal facility, mice were housed in groups of 3 for 2 weeks in standard sized cages. At 9 weeks of age, groups of 12 mice were weighed and placed into large, structurally complex, custom built vivaria (length 150cm, height 80cm, width 80cm; Mid-Atlantic; **Figure S6**). The vivaria were constructed as described in Williamson et al.^26^. Each vivarium consists of an upper level constructed of multiple shelves connected by plastic tubes and covered in pine bedding and a lower level comprised of 5 interconnected standard sized cages filled with pine bedding and connected by a system of plastic tubes. Mice can access all levels of the vivarium at any time through this interconnecting system of ramps and tunnels. Standard chow and water were provided ad libitum on the top level of the vivarium. Social groups were created such that in each group of 12 females, each individual had previous social experience with maximum only one other individual and at least 6 females per group had absolutely no experience with any of the other individuals. Mice were placed in the vivarium at the onset of the dark cycle on Day 1 of the experiment and were observed by trained observers for 2 hours directly following introduction to the group and for 2 hours each day for the next two weeks (Day 1 – Day 14). Observations always occurred during the dark cycle at some point during the first 6 hours of lights off (red light). During these live observations, observers used all occurrence sampling to record the winner and loser in all instances of fighting, chasing, mounting, subordinate posture, and induced-flee behaviors (see **Table S1** for an ethogram of these behaviors). Winners of each encounter were considered to be those that chased, bit, mounted, or forced another individual to exhibit subordinate behavior. If behaviors between two females co-occurred within 2 seconds of each other they were recorded with the priority fighting, chasing, mounting, subordinate posture, flee. This method has been used previously in our lab to understand the social organization of groups of male mice^5,6,26,27,95,96^. Vaginal smears were collected from every mouse each evening at the same time of day (six to eight hours post lights-off). To collect the samples, trained lab members removed mice from the vivaria individually and placed them back as soon as the sample was collected. Collecting samples from each social group interrupted the group for less than 5 minutes. Smear samples were analyzed under a microscope by a single trained lab member and double checked by a second lab member to verify accuracy. Mice were weighed, final estrus smears taken, and euthanized via decapitation 2 hours post lights off on Day 15. Trunk blood was collected into heparinized tubes, immediately placed on ice, centrifuged at 4°C in a refrigerated centrifuge, and plasma separated and frozen at −80°C until analyzed for corticosterone and estradiol levels via radioimmunoassay. Brains were collected and flash frozen in hexane and stored at - 80°C until dissection. At the end of group housing, the 2 most dominant and 2 most subordinate individuals were determined using the Glicko Rating System^26,97^ as well as David’s Scores^26,98^. Plasma hormone and brain mRNA levels were measured for these two most dominant and two most subordinate individuals in each group, except for in two cohorts where it was difficult to distinguish the beta and gamma female so three dominant individuals and two subordinate individuals were used.

### Hormone assays

Plasma corticosterone and plasma estradiol concentrations were measured using commercially available kits (MP Biomedicals) and conducted using the manufacturer’s specifications. For the corticosterone assay, the average inter-assay coefficient of variation was 9.3%, the lowest detectable was 24.78 ng/ul, and the highest detectable was 938.34 ng/ul. For the estradiol assay, the coefficient of variation was 7.2%, the lowest detectable was 8.53 pg/ul, and the highest detectable was 2455.79 pg/ul.

### Gene expression

Brains were stored at −80° C until dissection. Samples of the medial preoptic area (mPOA) and ventromedial hypothalamus (VMH) were collected using a Harris Micro-Punch with reference to the coronal plane from the Mouse Brain Atlas^99^ and the Allen Brain Atlas^100^. The mPOA was collected as one 1mm diameter area along the midline from Bregma +0.14 mm to −0.7 mm. The VMH was collected as one 1mm diameter area from each hemisphere from Bregma −1.34 mm to −1.82mm. RNA was isolated from both brain regions using the AllPrep RNA Micro Kit (Qiagen) and reverse transcribed to cDNA using the SuperScript III First-Strand Synthesis System for RT-PCR applications. Quantitative RT-PCR was performed with 1ul of cDNA using an ABI 7500 Fast Thermal Cycler and the Fast SYBR Green Master Mix reagent (Applied Biosystems). All primer probes (Sigma-Aldrich) were designed to span exon boundaries ensuring amplification of only mRNA. The following validated quantitative PCR primers were used for mRNA analysis: estrogen receptor alpha (ERα – Forward: CGTGTGCAATGACTATGCCTCT; Reverse: TGGTGCATTGGTTTGTAGCTGG), estrogen receptor beta (ERβ – Forward: GTCAGGCACATCAGTAACAAGGG; Reverse: ATTCAGCATCTCCAGCAGCAGGTC), progesterone receptor (PR – Forward: GCGAGAGACAACTGCTTTCAGT; Reverse: CAAACACCATCAGGCTCATCCA), gonadotropin releasing hormone (GnRH – Forward: AGCACTGGTCCTATGGGTTG; Reverse: GGTTCTGCCATTTGATCCAC), oxytocin receptor (OTR – Forward: TTCTTCGTGCAGATGTGGAG; Reverse: CCAAGAGCATGGCAATGATG), opioid receptor µ 1 (OPRM1 – Forward: AATGTTCATGGCAACCACAA; Reverse: TTTGAGCAGGTTCTCCCAGT).

### Statistical analysis

All statistical analyses were undertaken in R v.3.5.0^101^.

### Group dominance structure and social organization

For each cohort, six measures of dominance structure and organization were calculated: Landau’s modified *h’*, directional consistency, steepness, triangle transitivity, despotism and Gini’s coefficient of wins and losses. The methods of calculation for these measures are detailed in Williamson et al.^26^, but briefly: Landau’s modified *h’*, directional consistency, and steepness are calculated using frequency win/loss sociomatrices, which are created using the total frequency of wins and losses recorded for each individual over the observation period. Landau’s modified h’ evaluates the extent to which individuals in a hierarchy can be linearly ordered^98^ and ranges from 0-1, with a value of 1 indicating a completely linear hierarchy. Triangle transitivity measures the proportion of relations between all triads that are transitive (i.e. if individual A is dominant to individual B and individual B is dominant to individual C, then individual A is dominant to individual C; a perfect hierarchy would have all transitive triads)^102^. It is calculated using a binary win/loss sociomatrix, where 1s are assigned to individuals in rows that won more often against individuals in columns and 0s are assigned to individuals in rows that lost more often to individuals in columns. Triangle transitivity ranges from 0-1, with 1 indicating that all triads are transitive (i.e. a perfectly linear hierarchy). Steepness measures the unevenness of relative individual dominance within the hierarchy. This is calculated from the relative distribution of David’s Scores, a win proportion measure adjusted for strength of opponents^103^. It ranges from 0-1 with a score closer to 1 indicating that power is not equitably distributed across the hierarchy, but rather lies in the hands of a few powerful individuals at the top. Directional consistency measures the degree to which all agonistic interactions occur in the direction from the more dominant to more subordinate individual in the pair. It also ranges from 0-1, with 1 indicating that all agonistic interactions occur in the direction of dominant to subordinate. Significance testing for Landau’s modified *h’*, triangle transitivity, steepness and directional consistency were carried out using appropriate randomization methods^26^. P-values represent the proportion of times that values were observed in randomized data that were greater than or equal to the observed values from empirical data. Despotism is the proportion of all wins by the dominant male over the total number of aggressive interactions over the observation period. It is a value between 0-1, with 1 indicating that the alpha male performed 100% of all aggression within the group. Gini-coefficient is a measure of equality versus inequality in a distribution. We calculated the Gini-coefficients for the frequency of wins and losses by each animal across cohorts. It ranges from 0-1. Values closer to 1 indicate more inequality meaning that a higher number of wins/losses are associated with relatively few individuals. Values closer to 0 indicate that the frequency of wins/losses are equally distributed across all individuals. We examined the association between initial body weight on Day 1 of group housing and final social rank for each social group using Spearman Rank correlation tests.

Landau’s modified *h’*, triangle transitivity, directional consistency and despotism were calculated using the R package ‘compete’ v.0.1^104^. Steepness was calculated using the R package ‘steepness’ v.0.2.2^105^. Gini coefficients were calculated using the ‘ineq’^106^ package.

### Comparison between fighting, chasing and mounting behaviors

Raw frequency sociomatrices were constructed for each cohort based separately on fighting, chasing and mounting behaviors. To determine how correlated each matrix (fighting, chasing or mounting) was to each other within each cohort we performed a Quadratic Assignment Procedure (QAP) test with 1000 Monte Carlo randomizations of the data using the ‘sna’ package v.2.4^107^. From these matrices the directional consistency and Gini-coefficient of wins and losses was also calculated for each behavior for each cohort. To test if differences in these values existed between behaviors across cohorts we used Friedman Tests. Significant differences between behaviors were determined using Nemenyi Post-Hoc Tests using the ‘PMCMR’ R package v.4.3^108^.

### Emergence of hierarchies and individual ranks

To determine how quickly each cohort formed a linear hierarchy we calculated Landau’s modified *h’* and triangle transitivity as well as running significance tests for the aggregated win-loss data for each cohort up to each day. The emergence of individual ranks across time was identified using Glicko ratings. All individuals begin with a Glicko rating of 2200, and points are added or subtracted based on winning or losing against other individuals. The degree of points won or lost is dependent upon the difference in ratings between the two individuals. If an individual with a high Glicko rating defeats an individual with a low Glicko rating, relatively few points would be added to their total and relatively few points would be subtracted from the defeated individual. If an individual with a low Glicko rating defeats an individual with a high Glicko rating, a larger number of points would be added to their total and subtracted from the loser. Glicko ratings of all individuals in each group are recalculated after every behavioral interaction. We used a constant value of 3 in our calculations of Glicko ratings. See Williamson et al.^26^ for more information. Glicko ratings were calculated using the ‘PlayerRatings’ package v.1.0 in R^109^.

### Comparison of female social hierarchy behavior to male social hierarchy behavior

To measure differences in social hierarchy structure between male and female social groups, we compared female data from this study with previously published data on male social hierarchies from Williamson et al.^26^. We recalculated hierarchy measures using the first fourteen days of observation data from the 10 groups of 12 male CD-1 mice who were housed and observed in exactly the same manner as the female groups in the current study. We used Wilcoxon rank sum tests to compare the values of Landau’s modified h’, triangle transitivity, steepness, directional consistency, despotism and Gini-coefficient of wins and losses between male and female groups.

### Analysis of hourly occurrence rate of each agonistic behavior

All Bayesian linear and generalized linear regressions were fitted using R package ‘brms’^110^. For each fighting, chasing, and mounting behavior observed throughout the group housing period for each cohort, we tested whether the day of group housing affects hourly occurrence rate of each behavior by fitting the data with a gaussian distribution. Then we analyzed if the hourly occurrence rate of each behavior differs by individual final social rank using a two-process hurdle-gamma family. In this model, the probability of having zeros as the occurrence rates are modelled with a binomial error distribution with logit link function and non-zero non-integer continuous values are fitted with a gamma error distribution with log link^111^. We chose to use this model as the hourly occurrence rate data contains a significant number of zeros as subordinate individuals often barely initiate agonistic behaviors and alpha individuals barely receive aggression especially once the hierarchies are established. Further in this model we assume the difference of any effect of individual ranks is not necessarily equidistant between ranks. We therefore treated individual social rank as a monotonic predictor rather than a continuous variable throughout the entire statistical analysis^112^. The beta coefficients estimated from monotonic models presented in this study indicate the direction and the size (range between the lowest and highest categories of the ordinal fixed factor e.g. rank) of the effects.

### Analysis of estrus cycle state

We tested whether a mouse with higher social rank in the hierarchy is more likely to be in one estrus state compared to another by fitting a multinomial logistic mixed effect model with estrus status as the outcome variable, individual social rank as a monotonic predictor and cohort ID and subject ID as random effects. We tested whether an individual is more likely to give (hourly given aggression rate) or receive (hourly received aggression rate) each agonistic behavior when the mouse is in a certain estrus status using a hurdle-gamma model. First, the probability of the hourly occurrence rate being zero was predicted for each individual with daily Glicko rank of each mouse as a monotonic predictor to test whether the observed number of zeros in the data could be explained by rank. After this hurdle, the non-zero values of the hourly occurrence rate were fitted with a predictor of estrus state for each individual across each group housing day. Cohort ID, subject ID and day of group housing were set as random effects for both processes.

### Analysis of plasma corticosterone and estradiol levels

Using leave-one-out cross-validation information criteria (LOOIC)^113^, we compared four models with different combination of fixed effects: i) dominant-subordinate status, ii) last measured estrus cycle before blood collection, iii) dominant-subordinate status and last measured estrus state, iv) the interaction of social status and estrus state. All models were fitted with cohort ID as a random effect. For corticosterone data, a model with dominant-subordinate status only resulted in the best fit. A model with the status and the estrus state resulted in the best fit for estradiol data.

### Analysis of gene expression levels

Using the R package ‘MCMC.qpcr’^114^, we analyzed the differences in gene expression between dominant and subordinate mice by fitting a generalized mixed effect model with Poisson-lognormal distribution and a Bayesian Markov Chain Monte Carlo sampling approach. This approach provides advantages compared to standard delta-CT analysis as it accounts for random variation between duplicates, increases power by analyzing data for all target genes in one model, and does not require control genes^115^. Briefly, the raw threshold cycle (CT) values were converted into molecule count data with consideration of the amplification efficiency of each gene. We fitted the model with dominant-subordinate status category as a fixed factor and cohort ID and subject ID as random factors. We confirmed linearity of the model by inspecting diagnostic plots.

### Data availability statement

All raw data and code used in this paper are publicly available at GitHub https://github.com/jalapic/females

## Acknowledgements

We thank Dr. Frances Champagne for advice and suggestions in writing the manuscript and Curley Lab students for help with behavioral observations.

## Author contributions statement

CMW, WL, ARD and JPC conceived and planned the experiments. CMW, WL, ARD, RDR and AL carried out the behavioral work and husbandry. RDR planned and carried out the hormone analysis. CMW and ARD carried out the gene expression experiment. CMW, WL and JPC analyzed the data. CMW, WL, & JPC wrote the paper. All authors provided critical feedback and helped shape the research, analysis and manuscript.

## Additional information

### Competing interests

The authors declare there are no competing interests.

