## Supplemental tables and figures for "Social hierarchy position in female mice is associated with plasma corticosterone levels and hypothalamic gene expression"

**Supplementary Figure S1.** Binarized sociomatrices of wins and losses for each cohort. A 1 in a cell represents that the individual in the row won more fights than they lost against the individual in the column. The degree of redness represents the directional consistency of wins and losses for each dyad. Individuals are ordered by I&SI rank.

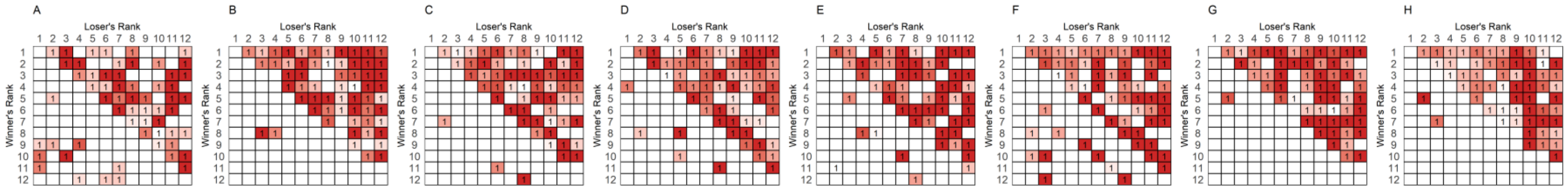

**Supplementary Figure S2.** Boxplots showing the total number of fighting, chasing and mounting events per hour of observation by rank across the fourteen days of observation.

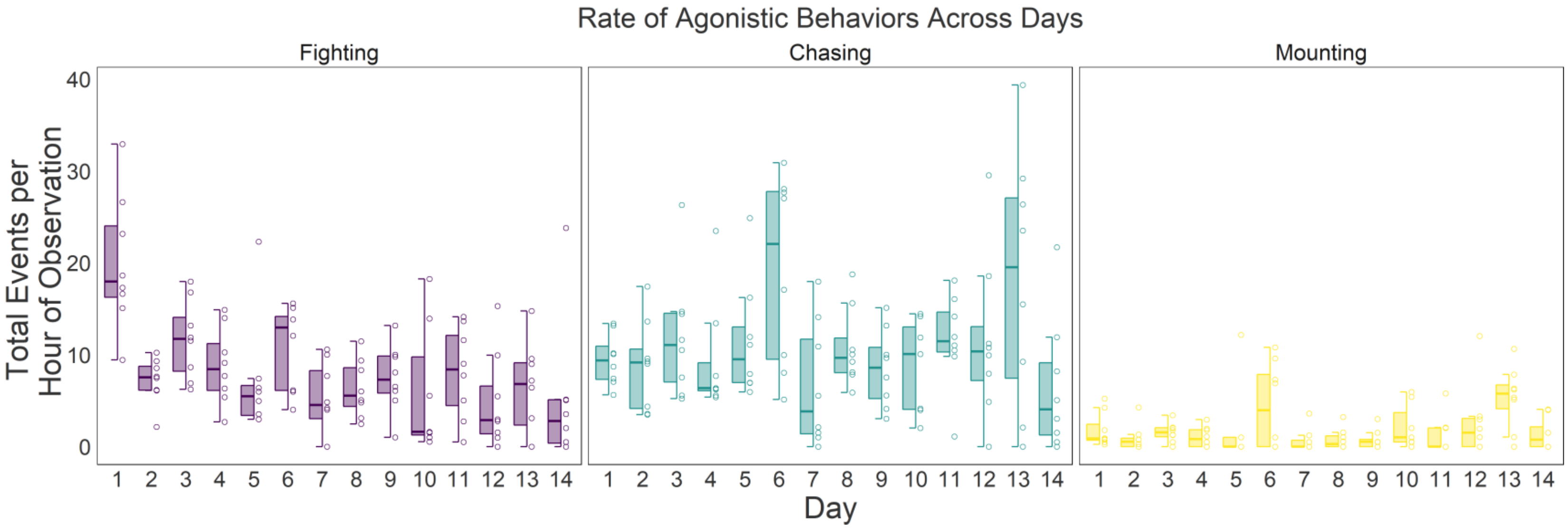

**Supplementary Figure S3.** Raw sociomatrices of wins and losses for each behavior across all cohort (A-H). Row 1 = fighting; Row 2 = chasing; Row 3 = mounting. Each value represents the total number of wins by the individual in each row against the individual in each column for each behavior. The degree of redness represents the frequency of wins. Individuals are ordered by I&SI rank.

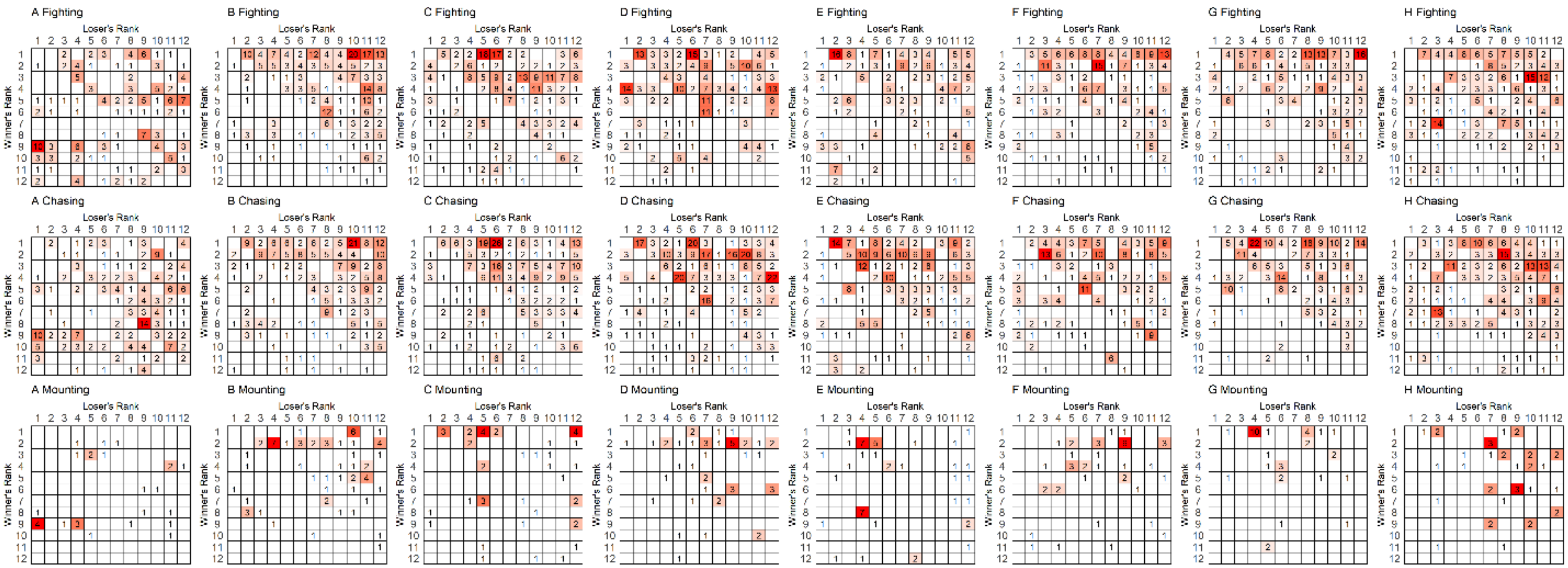

**Supplementary Figure S4.** The directional consistency (DC) and Gini-Coefficient of Wins and Losses of each behavior (fighting, chasing, mounting). Mounting behavior is significantly more directionally consistent and has a higher Gini-Coefficient of wins and losses than other behaviors.

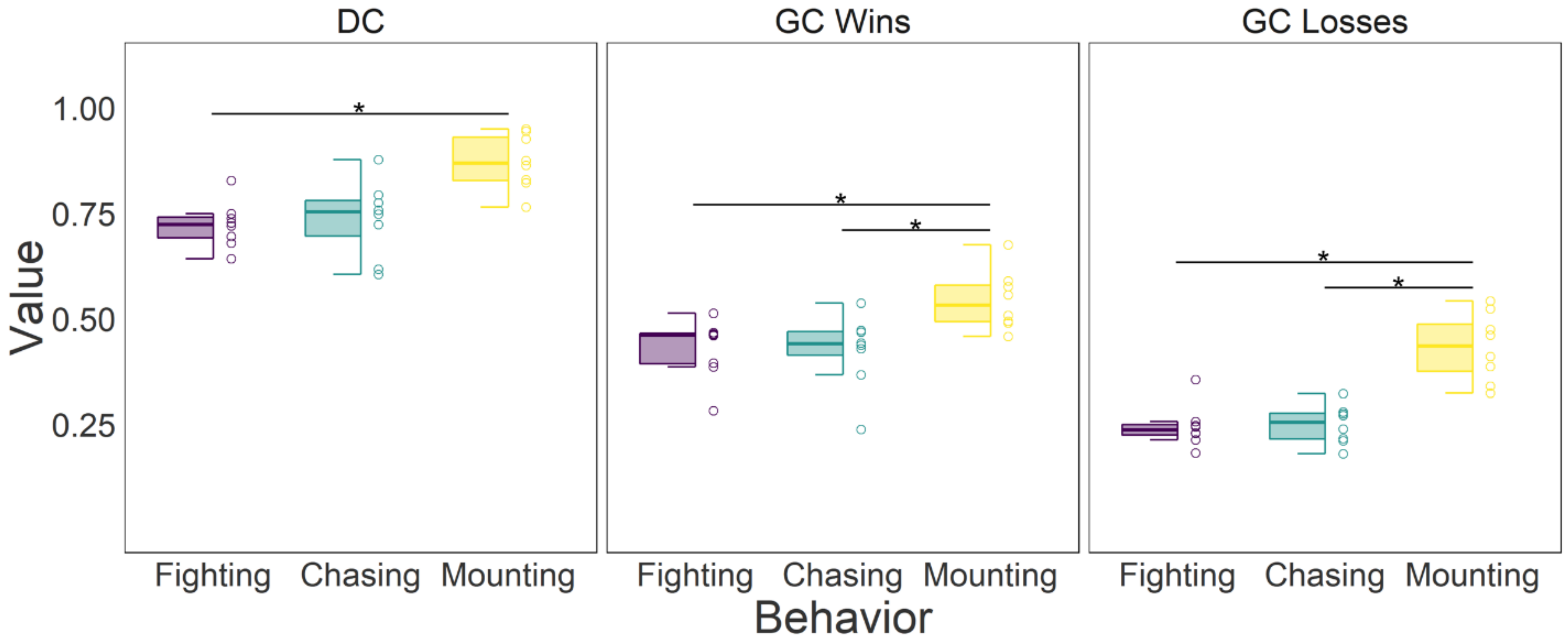

**Supplementary Figure S5.** The proportion of all wins that are fights (purple), chases (green) or mounts (yellow) by rank across all cohorts. There is no effect of rank on the proportion of each behavior used.

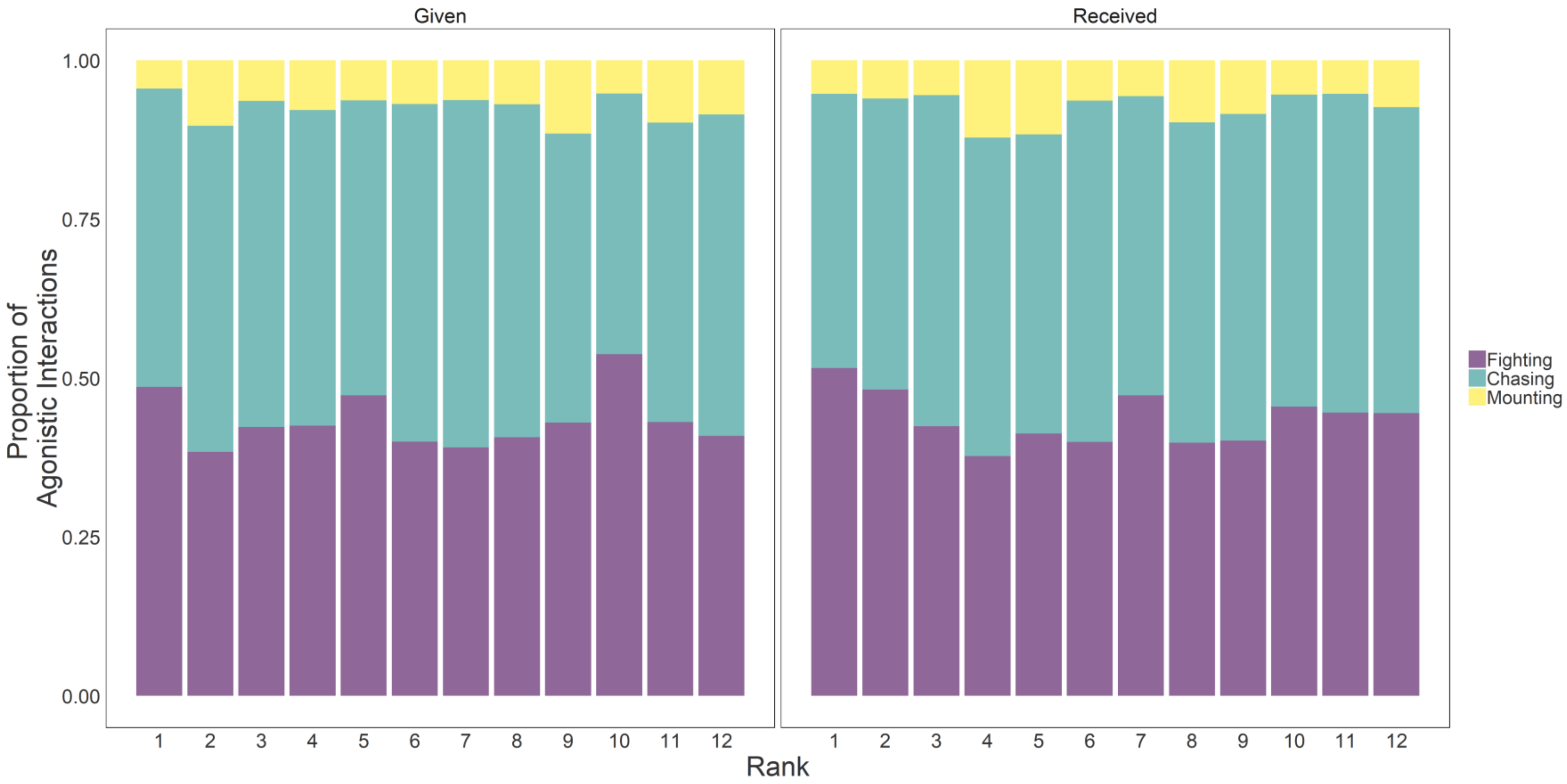

**Supplementary figure S6.** Vivarium for social group housing. The vivaria consists of two levels. An upper level comprised of ramps, shelves, ladders and physical enrichment. Food and water are available *ad libitum* at the top of the unit. The bottom level is accessible via two tunnels from the floor of the top level. Tubes interweave between five nest boxes. A red plastic sheet (not shown in picture) surrounds the lower level to ensure darkness of this level throughout housing.

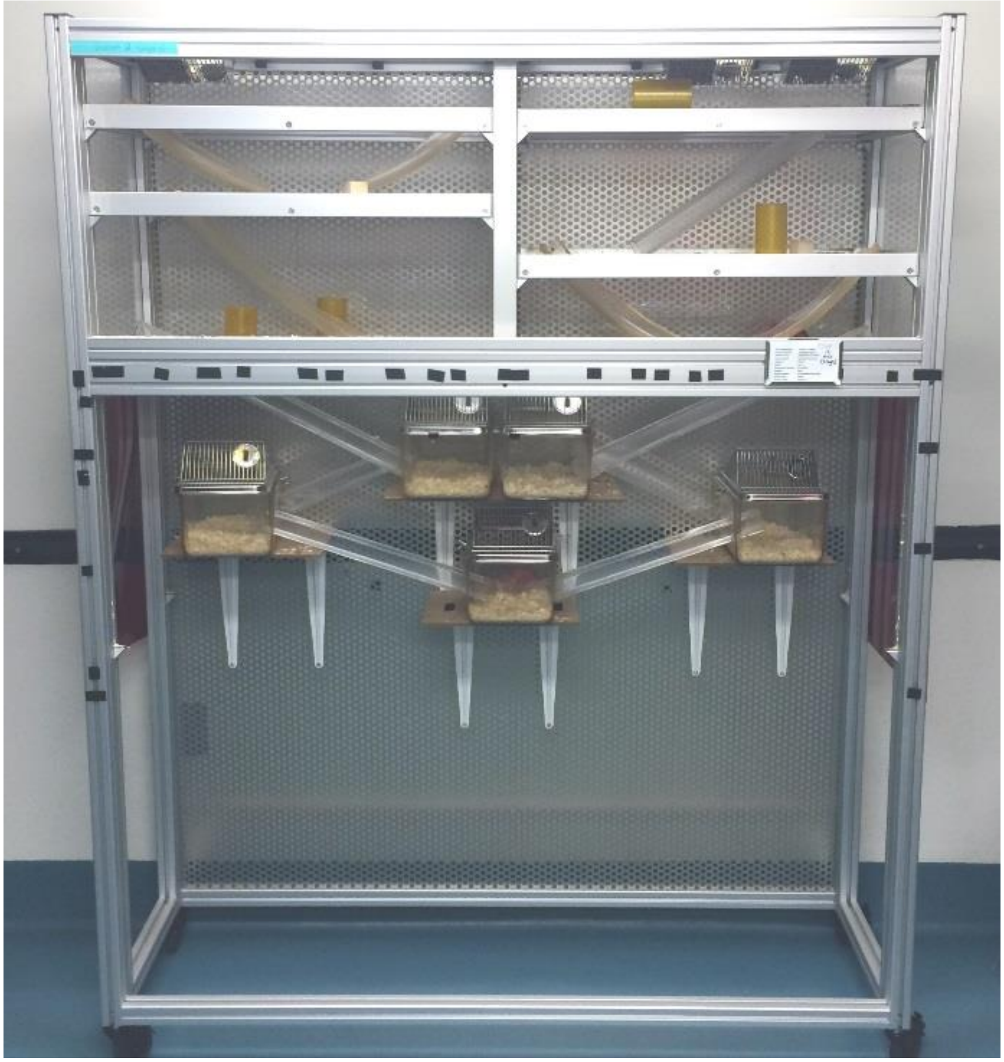

**Supplemental Table S1.** Mouse social behavior ethogram.

| Priority | Behavior | Description |
|---|---|---|
| 1 | Fighting | Individual lunges at and/or bites the other individual |
| 2 | Chasing | Individual follows the target individual rapidly and aggressively while the other individual attempts to flee |
| 3 | Mounting | Individual mounts another individual from behind with the recipient attempting to flee or otherwise being pinned to the floor |
| 4 | Subordinate posture | Individual responds to the approach from another individual by remaining motionless and/or exposing their nape |
| 5 | Induced-flee | Individual flees without any aggression shown by another individual |
